# Morphometric Interpretation of Postganglionic Sympathetic Neurons that Innervate Myocardium

**DOI:** 10.1101/2025.07.27.667010

**Authors:** Sachin Sharma

## Abstract

**Background and Aims:** The sympathetic nervous system modulates cardiac functions through neurotransmitters such as neuropeptide-Y and galanin released by postganglionic neurons. Hence, we hypothesize that dendritic and axonal morphological architecture of cardiac-innervating neurons might reflect the communicating input and output signals either within the postsynaptic neurons or to the adjacent myocardial cells.

**Methods:** In the current study, we carried-out morphometric analyses of cardiac-innervating neurons, measuring dendritic size, shape, and neuronal-polarity. We used retrograde tracers (adeno-associated virus conjugate to fluorophores) injected either from left-ventricular myocardial tissue or fore-limb skin tissue-beds. Stellate ganglia were harvested from the mice for imaging and morphometric analysis.

**Results:** Our findings revealed that cardiac-projecting neurons exhibit a multipolar structure and are significantly larger in cross-sectional area and volume compared to forelimb skinpad-innervating neurons.

**Interpretation:** These morphological characteristics may offer valuable insights into the neural architecture underlying cardiac remodelling, although further investigation is needed. This study focuses solely on the structural, not functional, features of cardiac-innervating neurons to better understand their specialization within the autonomic nervous system.

## Introduction

The clinical interest in autonomic nervous systems lies within the extra cardiac as well as intrinsic cardiac nervous system [1]. Cardiac-neuroaxis has been under-investigation for several decades with scant attention being paid to morphological features of the myocardium-innervating neurons [2], [3]. The soma (cell-bodies) of postganglionic sympathetic neurons (PGNs) primarily resides within the cervicothoracic sympathetic paravertebral chain particularly stellate ganglia and middle cervical ganglia and, innervates ventricles of the myocardium uniformly [4], [5], [6]. The development and regulatory mechanism of sympathetic innervation of myocardial tissues remain elusive, as does the role of this innervation in cardiomyopathy and/or arrhythmogenesis. Several studies reported release of neurochemicals such as Sema3a, endothelin, and neurotrophic growth factor play an important role in homogeneous sympathetic innervation while the release of catecholamines, neuropeptide-Y, and Galanin act on myocardial tissues and regulate the cardiac output functions [7], [8], [9], [10], [11].

Neurons, a polarized cell with a distinct dendritic and axonal territories, encompassing cell membrane that fine-tunes numerous types of signals, propagate them in the form of electrical pulses [12]. The dendritic and axonal morphologies within a neuron are the extended plasma membrane protrusions that reflect as communicating medium for signals and considered as the defining characteristics of specific neuronal subpopulations. Dendrites exist in various shapes and sizes and, their morphologies have been used for neuronal classification [13]. Dendritic structures influence the number of synapses and wiring logic within the neuronal networks and secondly, it impacts spatial and temporal integration of postsynaptic potentials [14], [15]. Neurons with bipolar dendritic morphology are more receptive in regulating their input-output relationship [16]. Dendrites and axons, the beautiful arbours of neurons, transmit information back to their synaptic inputs by releasing the neuroactive substances, primarily neuropeptides as evidenced by their release in the brain [17]. The hypothalamic dendritic neuropeptides and somatodendritic dopamine have significant physiological impact on brain function and maintenance [18]. These observations suggest that dendritic and axonal surfaces play important roles in both signal transmission and the exchange of molecules with the surrounding environment [12].

In this study, we investigate the morphology and structural characteristics of cardiac-projecting neurons located in the stellate ganglia, aiming to explore potential relationships between their morphological features and functional roles.

## Methods

### Animals

Mice strain C57BL/6J (000664) and *NPY-*hrGFP (006417) were purchased from Jackson Laboratories. The study (protocol number: **18–048**) was approved by the UCLA Institutional Animal Care and Use Committee. The ethical approvals for the use of adeno-associated virus vectors (AAVs) were provided by the Institutional Biosafety Committee (IBC), UCLA. Adult male mice, aged between 8 to 10 weeks, were housed according to the standard laboratory conditions (12 hr light/dark) with ad libitum access to food and water.

### AAV injections and retrograde tracing

#### Ventricle injections

Mice were pretreated with carprofen (5 mg/kg, s.c.), cefazolin (Hikma Farmaceutica, #01439924-90), and buprenorphine (0.05 mg/kg, s.c.) one hour prior to surgery. Animals were anesthetized with isoflurane (induction at 4%, maintenance at 2%, inhalation), intubated, and mechanically ventilated. Core body temperature was measured and maintained at 36°C. The surgical incision site was cleaned three times with 100% povidone iodine and 70% ethanol in H_2_0 (vol/vol). A left lateral thoracotomy was performed at the fourth intercostal space, the pericardium opened, and ventricles were exposed. 10μl of AAV conjugated to GFP (1e+12 gc/ml) (Viral vector core, Boston Children’s Hospital) were administered intramyocardially in left ventricle with a 31-gauge needle. The surgical wounds were closed with 6–0 sutures. Buprenorphine (0.05 mg kg-1, s.c.) and cefazolin (Hikma Farmaceutica, #0143-9924-90) were administered once daily for up to three days post-surgery. Animals were sacrificed, perfused with 4% paraformaldehyde (4% PFA), and tissues were harvested 3-weeks postsurgery of AAV injections.

#### Forelimb injections

Mice were anesthetized with isoflurane (Induction at 5%, maintenance at 1-3%, inhalation) and forelimb skin-pads were cleaned 3 times with 70% ethanol in H_2_O (vol/vol). 10μl of AAV conjugated to tdTomato (1e+12 gc/ml) (Viral vector core, Boston Children’s Hospital) were injected on forelimb tissue-pads. Three weeks post-surgery, animals were sacrificed, perfused with 4% PFA and paravertebral sympathetic chain ganglia (right and left stellate ganglia) were isolated.

#### Stellate ganglia isolation

To isolate stellate ganglia from AAVs injected in heart and forelimb from three weeks of postsurgery, mice were anesthetized with 3% isoflurane for 5 min until no reflexes were observed. Afterwards, cervical dislocation was carried out and mice were perfused with 50 mL ice-cold 0.01M phosphate buffer saline (PBS) (10 mM potassium phosphate, pH 7.4 and 150 mM NaCl) containing heparin followed by 50 mL freshly prepared, ice-cold 4% paraformaldehyde in PBS. Once mice were fixed, stellate ganglionic tissues were isolated using a dissecting microscope. Isolated tissues were fixed in 4% PFA overnight at 4°C, washed, and stored in 0.01M PBS at 4°C. Stellate ganglia were collected bilaterally (both right and left sides) from all animals. For whole-mount preparations, tissues were mounted on the slides under light-protecting sheet.

#### Confocal microscopy and image processing

High-fluorescent images were acquired on a confocal laser scanning microscope LSM 880 Upright (Carl-Zeiss, Germany). The laser lines used were 405 nm (diode [405-30]) for excitation of DAPI, 488 nm (argon) for Alexa-488, 561 nm (DPSS [561-10]) for Cy3, and 633 nm (HeNe) for Alexa-647. Fluorescence and bright-field illumination modes were used during the image acquisition process. Samples were visualized through Plan-apochromat 10x/0.45 air, Plan-Apochromat 20x/0.8 air, and Plan-Apochromat 63x/1.4 oil objectives. Each fluorophore was used in a separate cohort of animals to avoid cross-contamination. No dual labeling was performed in the same animal. The fluorophore choice allowed for better separation in confocal imaging but was not intended for simultaneous visualization in vivo.

For imaging the whole-mount tissues, we used z-stack optical sectioning and tile-scan image overview mode to generate the composite images. High-resolution images of individual labelled neurons including axons and dendrites were analysed using Imaris software. We used ‘statistic’ Imaris tool to measure the volume and cross-sectional surface area from soma region of the individual cardiac and forelimb innervating neurons. The image analysis software ImageJ was used to count the number of projections, long-axis length, cross-sectional area, and volume. During analysis, we found no significant morphological differences between neurons labeled in the left versus right stellate ganglia. Therefore, data from both sides were pooled for analysis. All our morphometric analysis for cardiac- and forelimb-injected mice was conducted on control (C57BL/6J) mice strain. We included only those neurons that were completely filled, optically isolated from neighbouring cells after thresholding, and showed clear and intact dendritic/axonal structures in 3D reconstructions. We used NPY-hrGFP mice only for the whole mount illustration of the stellate ganglia (Figure 2a).

#### Statistical analyses

GraphPad Prism 9.0.1 was used for data handling, analysis, and graph generation. Statistical tests performed are indicated in the figure-legend for each figure. Small sample size is a constrain in the study, hence, we did not use statistical tests to assess the normality of datasets.

For statistical significance, we performed Mann-Whitney non-parametric test to analyse the data. All data are shown here as individual data points. We provided exact (*p-value*) value for each experiment in figure legends and indications of *p*-values are shown as *=*p*<0.05, **=*p*<0.01, ***=*p*<0.001, ****=*p*<0.0001 in figures.

## Results

In this study, we examined the morphological features of cardiacversus forelimbinnervating neurons using retrograde viral tracers. We characterized neuronal structural polarity, neuronal length, cross-sectional area, and volume in cardiac-projecting neurons and compared them with those innervating the forelimb. The results are presented in the following order:

### Neuronal structural polarity

Cell polarization is a critical process in various cellular functions, including cell differentiation, morphogenesis, and migration. Neurons, being highly polarized cells, typically develop a single axon and multiple dendrites, forming cellular compartments of protein content and electrical properties.

Characterization of individual neurons is an essential for understanding the functioning of how neural circuitry operates. The AAV serotypes used in our study are retrogradely transported and predominantly label neurons projecting to the injection site. Although we did not use transgenic reporters to define neurotransmitter identity in this study, our prior publication extensively characterizes the neurotransmitter diversity in stellate ganglia [19]. Recent anatomical studies have also revealed the presence of nitrergic, non-parasympathetic cardiac fibers, further expanding the known diversity of cardiac-projecting neurons [20]. Using high-resolution confocal microscopy, we visualized cardiac-projecting (**GFP**^**+**^) and forelimb-innervating (**tdTomato**^**+**^) neurons in stellate ganglia (Figure 1 and SF 1). Figure 1(ai) presents a cardiac-innervating neuron expressing GFP in somata as well in dendrites. The image was a microscopic image while adjacent figure 1(**aii**) displays an *Imaris* generated image of the same microscopic image. Similarly, figure 1(**aiii**) shows a forelimb-innervated multipolar neuron with expression of tdTomato filled across soma and dendrites while adjacent figure 1(**aiv**) represents its corresponding *Imaris* generated image. Figure 1(**bi**) and (**bii**) depict a pseudo-unipolar neuron labelled with retrograde tracers in both microscopic and Imaris generated images, respectively. Figure 1(**ci**) and (**cii**) illustrate a unipolar neuron labelled with retrograde tracers, shown in microscopic and *Imaris*-generated formats.

**Figure 1.**
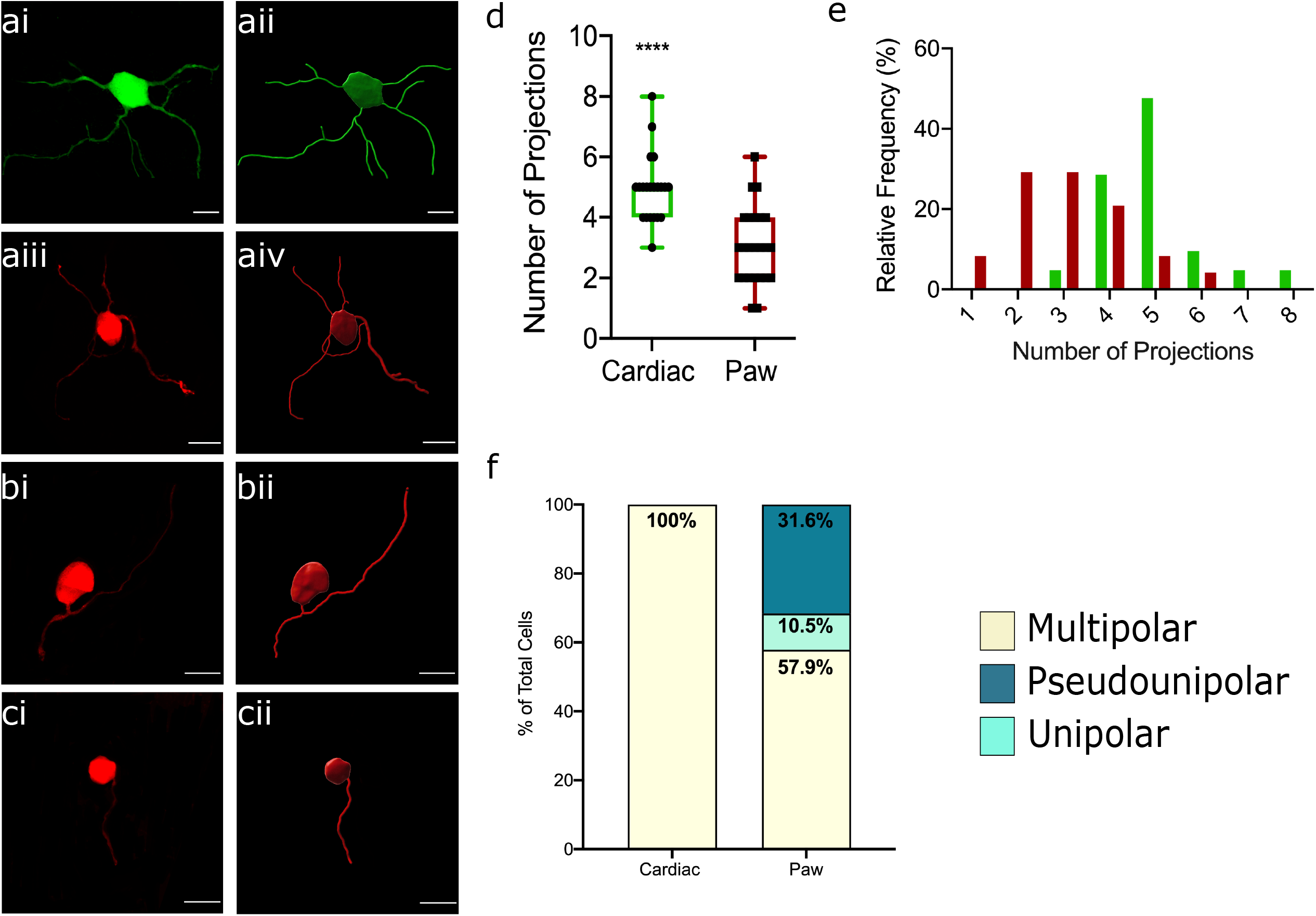
Morphometric measurements of the cardiac and forelimb neurons. Paired images of (**a**i) cardiac (green) and (**aii, bi, ci**), forelimb (red) neurons are shown. The right face of row in (**ai, aiii, bi**, and **ci**) represent the original microscopic image and left side of row (**aii, aiv, bii**, and **cii**) has intensity-based reconstruction images by Imaris from the original images as shown in right side of the row. **d**, Comparison of numbers of projections (p<0.0001). **e**, Histogram representing projections for cardiac neurons (sample size = 21) and forelimb (sample size = 25) neurons. **f**, “Parts of whole” demonstrated the share of morphological types of cardiac and forelimb neurons. (n=3 mice per group). Mann-Whitney non-parametric test was used to analyse the data. *Scale bars: 25 μm*

We counted dendrites-projections from cardiac-projecting neurons and compared from forelimb-innervating neurons. We found that the number of projections from cardiac-neurons were significantly higher compared to forelimb-innervating neurons (Figure **1d**). We also report the histogram of all cardiac- and forelimb-innervating neurons, representing the relative frequency and number of their projections (Figure **1e**). Our results indicate that all cardiac-projecting neurons were multipolar vs. forelimb-innervating neurons (Figure **1f**). In contrast, forelimb-innervating neurons 57% multipolar, 31.6% pseudo-unipolar, and 10.5% were unipolar in shape (Figure **1f**).

### Neuronal length, cross-sectional area and volume

The neuro-morphological features of a neurons are vital in their functional reorganization within tissues [21]. Neuronal length and surface area together correlate with number of synapses in dendrites and axon terminals [22], [23].

Figure 2(**a**) represents a tile-scan image of right stellate ganglia (RSG) from whole-mount preparations dual labelled with GFP (*Npy*-hrGFP) and tdTomato (forelimb-innervating neurons). High GFP expressing neurons are clustered within the cardiac-projecting region of stellate ganglia [6], [24]. Further to our investigation, we measured long-axis length, crosssectional area, and volume of cardiac- and forelimb-innervating neurons. We observed that long-axis length of cardiac-projecting neurons was significantly higher compared to forelimb-innervating neurons (Figure 2**b**). We observed similar patterns for the cross-sectional area (Figure 2**c**) and volume (Figure 2**d**) for cardiac-innervating neurons vs. forelimb-projecting neurons. Adjacent to each morphometric parameter of neurons, we presented histogram to show the relative frequency of respective neurons with their morphological features. All corresponding data for these measurements is provided within the supplemental information (Supplemental table 1).

**Figure 2.**
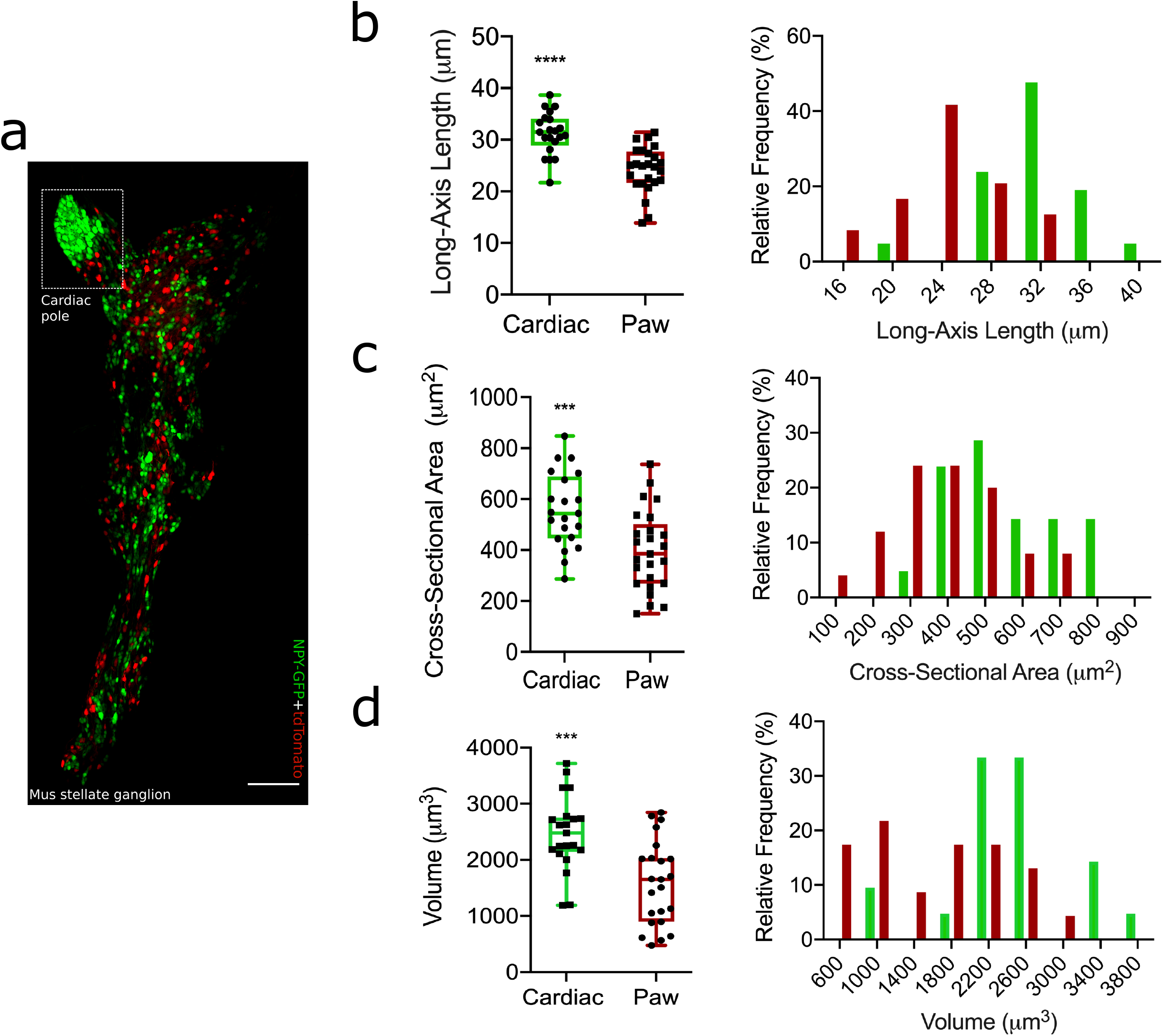
Morphometric measurements of the cardiac- and forelimb-innervating neurons. **a**, A tile-scan image of whole-mount right stellate ganglion isolated from AAV-tdTomato (red colored) injected in forelimb paw-pads of NPY-hrGFP (green) mouse strain. *Scale bar: 200 μm*. **b**, Long-axis length and histogram representation for cardiac neurons (sample size = 25) and forelimb neurons (sample size = 21), p<0.0001. **c**, Cross-sectional area and histogram representation for cardiac neurons (sample size = 21) and forelimb neurons (sample size = 25), p=0.0016. **d**, Volumetric analysis of cardiac (sample size = 21) and forelimb-neurons (sample size = 23) along with histogram representation, p=0.0003. n=3 mice per group. Mann-Whitney non-parametric test was used to analyse the data.

## 4. Discussion

Dendritic and axonal morphology are defining pattern of neuronal cell types and reflects upon their functional characteristics. To systemically analyse morphologies of cardiac-projecting neurons, we established a flow-chart/pipeline encompassing retrograde viral tracing, high-resolution imaging and image-reconstruction, and statistical computation using high-end software Imaris. We characterized number of projections, long-axis length, crosssectional area, and volume parameters for cardiac-innervating neurons vs. forelimb skin-pad innervating neurons for adult male mice.

Our study reveals that cardiac-innervating neurons differ significantly in morphology compared to forelimb-innervating neurons, particularly in neuronal structural polarity, dendritic complexity, and soma size. Therefore, we hypothesize that exclusive multipolar morphology of cardiac-projecting neurons aligns with their role in releasing/receiving multiple synaptic inputs in the form of neurotransmitters/neurochemicals, potentially enhancing their ability to modulate complex cardiac functions. These findings are being in consistent with previous studies suggesting that neuronal morphology correlates with functional integration in autonomic circuits [13]. Cardiac-innervating neurons exhibited significantly more dendritic projections than forelimb neurons, suggesting an increased capacity for synaptic integration. This enhanced dendritic complexity may facilitate the integration of various autonomic inputs, allowing cardiac-projecting neurons to regulate fine-tuned sympathetic outflow to the heart. Prior work has suggested that neurons with higher dendritic complexity exhibit greater synaptic plasticity and connectivity, potentially explaining why cardiac-innervating neurons demonstrate greater functional adaptability [15]. In a coronary-ligation induced infarct rabbit model, the neuronal enlargement of somata was observed in intrinsic cardiac innervating system from heart-failure group compared to controls [25]. Sanjay Singh and his co-workers have reported that hypertrophic vagal postganglionic neurons in cardiac ganglia in failing human hearts and hypertensive rats [26]. These studies clearly suggest neuronal plasticity within cardiac neurons during heart failure and myocardial infarction. Moreover, the AAV serotypes used in our study are retrogradely transported and predominantly label neurons projecting to the injection site.

Although we did not use transgenic reporters to define neurotransmitter identity in this study, previous work has shown that sympathetic neurons, including those in the stellate ganglia, can exhibit cholinergic differentiation independent of their target tissue [27]. This emphasizes the importance of interpreting retrograde labeling data with an understanding of underlying molecular diversity [6]. The significantly larger soma sizes, cross-sectional areas, and volumes of cardiac-innervating neurons compared to forelimb neurons indicate that cardiac neurons require increased cellular machinery to sustain high levels of neurotransmitter release and excitability. The observed differences in soma volume and dendritic complexity may reflect enhanced metabolic and electrophysiological demands, which could be necessary for continuous cardiac autonomic control [4].

The morphology of postganglionic sympathetic neurons is shaped not only by target innervation patterns but also by developmental cues such as neurotrophins, semaphorins, and activity-dependent pruning. Alterations in these signaling pathways can profoundly affect dendritic complexity and soma size. Furthermore, disease states such as heart failure or myocardial infarction are known to induce neuroplastic changes in sympathetic circuits, including hypertrophy, sprouting, and shifts in neurotransmitter phenotype. Understanding these dynamic processes is essential for interpreting morphological findings in both healthy and pathological contexts.

While our findings reveal distinct morphological features of cardiac-innervating neurons, we did not perform electrophysiological or functional assays. Therefore, any interpretations regarding excitability or synaptic integration remain speculative and warrant future validation.

## 5. Limitation

Current study is limited to only adult male mice. We do not report any sex-differences mediated changes in our results. Our work includes only smaller sample size, therefore, only a limited number of pairwise comparisons were conducted and corrections for multiple comparisons were not applied. We acknowledge this as a methodological limitation and note that future studies involving larger datasets and multiple testing scenarios will incorporate appropriate correction methods (e.g., Bonferroni or FDR) to control for type I error. Finally, this study was conducted exclusively in adult male mice. Given the known sex-based differences in autonomic regulation and cardiac remodeling, the findings may not fully generalize to female animals. Future studies should investigate potential sex-specific morphological features of cardiac-innervating neurons.

## 6. Conclusion

This study offers important insights into the distinct morphological characteristics of cardiac-innervating neurons, emphasizing their specialized role in autonomic regulation. Through systematic analysis of neuronal structural polarity, dendritic complexity, and soma size, we show that cardiac-projecting neurons possess a unique architecture that may enhance their capacity for synaptic integration and neurotransmission. These findings contribute to a deeper under-standing of neuro-cardiac communication and may help inform new strategies for addressing autonomic dysfunction in cardiovascular diseases. Future studies should investigate how these structural adaptations influence cardiac function under both physiological and pathological conditions, with the potential to guide the development of novel therapeutic approaches.

## Figure legends

**SF 1:**
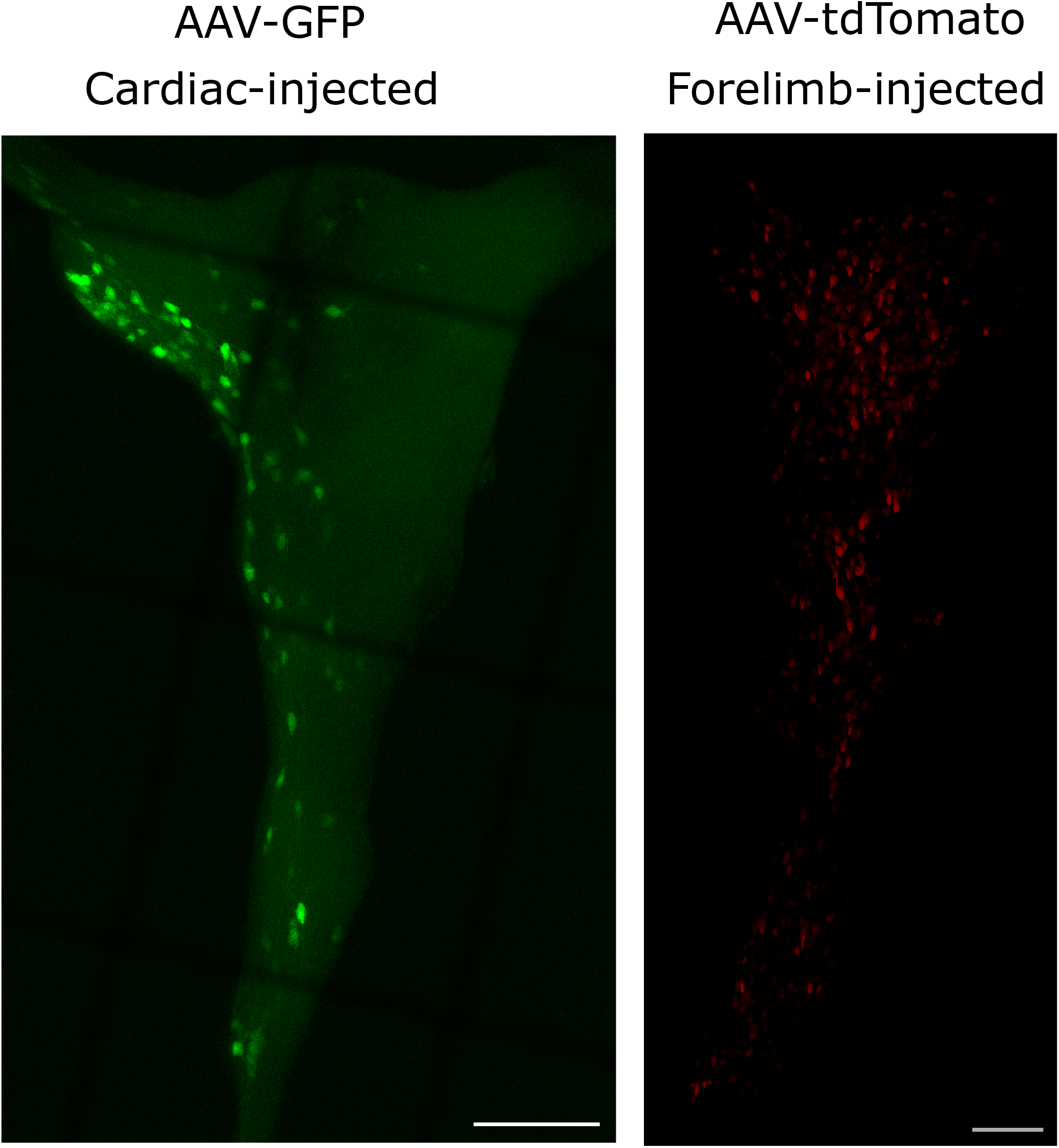
An image showing the visualization of cardiac-innervating (left) and forelimb-innervating (right) neurons within the stellate ganglia. The cardiac-injection was performed using AAV-GFP tracers while paw-injection was performed using AAV-tdTomato injections. *Scale bar: 200μm*

